# Increased intestinal permeability in an orally-reactive peanut allergy model identifies Angiopoietin like-4 as a biomarker

**DOI:** 10.1101/2021.07.14.452416

**Authors:** Erin C. Steinbach, Johanna M. Smeekens, Satyaki Roy, Takahiko Toyonaga, Caleb Cornaby, Layna Perini, Ana Berglind, Michael D. Kulis, Edwin H. Kim, Martin T. Ferris, Terrence S. Furey, A. Wesley Burks, Shehzad Z. Sheikh

## Abstract

Peanut allergy reaction severity correlates with increased intestinal epithelial cell (IEC) barrier permeability. CC027/GeniUnc mice develop peanut allergy by intragastric administration of peanut proteins without adjuvant. We report that peanut-allergic CC027/GeniUnc mice showed increased IEC barrier permeability and systemic peanut allergen Ara h 2 after challenge. Jejunal epithelial cell transcriptomics showed effects of peanut allergy on IEC proliferation, survival, and metabolism, and revealed IEC-predominant angiopoietin like-4 (Angptl4) as a unique feature of CC027/GeniUnc peanut allergy. CC027/GeniUnc mice and peanut-allergic pediatric patients demonstrated significantly higher serum Angptl4 and ANGPTL4 compared to control C3H/HeJ mice and non-peanut-allergic but atopic patients, respectively, highlighting its potential as a biomarker of peanut allergy.

## Introduction

Peanut is the number one cause of fatal food-associated anaphylaxis (1), and current management is limited to avoidance of peanut. Peanut allergy pathogenesis is incomplete, limiting development of new therapies. Evidence supports a critical role for epithelial cell (EC) dysfunction in peanut allergy. Conditions associated with genetic variants of the EC-specific barrier integrity genes *filaggrin* and *desmoglein-1* (2) and atopic dermatitis (3) are associated with increased risk of peanut allergy. The EC-specific cytokine thymic stromal lymphopoietin (TSLP) promotes allergic immune responses (4). Likewise, intestinal epithelial cell (IEC) barrier permeability is increased in patients with peanut allergy, which correlates with severity of reaction to peanut (5); greater serum peanut allergen levels are detected after ingestion in peanut-allergic compared to non-allergic patients (6). Factors that modify anaphylaxis severity (e.g., vigorous physical activity) are associated with increased IEC barrier permeability (7). Mechanisms underlying IEC barrier dysfunction in peanut allergy are poorly studied due to a lack of a robust animal model and limited access to patient samples.

Orgel, et al., previously identified a novel Collaborative Cross (8) mouse strain (CC027/GeniUnc, “CC027”) as susceptible to peanut allergy by intragastric (i.g.) administration without Th2-skewing adjuvant (i.e., *Cholera* toxin (CT)); another allergy-prone strain, C3H/HeJ (“C3H”), is not susceptible (9). Peanut-allergic CC027 mice produce peanut-specific IgE, accumulate intestinal mast cells, and have detectable serum peanut allergen (Ara h 2) after challenge (9). Circulating Ara h 2 suggests CC027 mice demonstrate IEC barrier dysfunction, thought this has not been studied. The CC027 mouse is a physiologically relevant model to study phenotypes and genomics that might better translate to humans than the current artificial models, which rely on non-physiological methods of sensitization or challenge, including i.v./i.p. sensitization with Th2-skewing adjuvants (alum, CT, staphylococcus enterotoxin B), large doses of allergen, and repeated oral challenges (10). CT induces a loss of IEC barrier integrity and supports an allergic immune response, potentially masking the independent effects of an allergen on IECs (10). Peanut itself has adjuvant properties (11), but its interaction with IECs has not been extensively studied.

We characterize IEC barrier dysfunction in the peanut-allergic CC027 mouse. Increased IEC barrier permeability and circulating Ara h 2 were associated with peanut allergy severity. Jejunal epithelial cell expression profiles in peanut-sensitized CC027 mice offered insights into IEC dysfunction. We identify angiopoietin-like protein 4 (*Angptl4*) as an IEC-enriched transcript that was significantly increased during peanut challenge and whose genomic location in the CC027 mouse is distinct from other mouse models of peanut challenge. Angptl4 is a secreted product (12); we report that serum Angptl4 levels were significantly elevated in peanut-sensitized CC027 mice, and that serum ANGPTL4 levels in peanut allergic pediatric patients were significantly higher than non-peanut allergic patients. This study highlights the CC027 mouse model as a unique tool to interrogate IEC barrier function and identify biomarkers in human peanut allergy.

## Results and Discussion

Peanut allergy was induced in female CC027 mice without adjuvant; a similar exposure protocol was applied to C3H mice as a control (**Figure 1A**) (9); CC027 but not C3H mice produced increased in peanut-specific IgE (13). Rectal temperatures, a surrogate of reaction severity, decreased significantly with time in peanut-sensitized & peanut-challenged (Peanut/Peanut) CC027 mice compared to PBS-sensitized & PBS-challenged (PBS/PBS) CC027 mice (**Fig. 1B**), but not in C3H mice (**Suppl. Fig. 1A**). PBS-sensitized & peanut-challenged (PBS/Peanut) CC027 mice demonstrated a transient temperature decrease 15 minutes after challenge that did not meet the threshold for a severe reaction (≥3°C decrease in rectal temperature) (**Fig. 1B**), suggesting that initial intestinal peanut exposure in CC027 mice leads to a transient mild reaction, perhaps facilitating sensitization.

**Figure 1.**
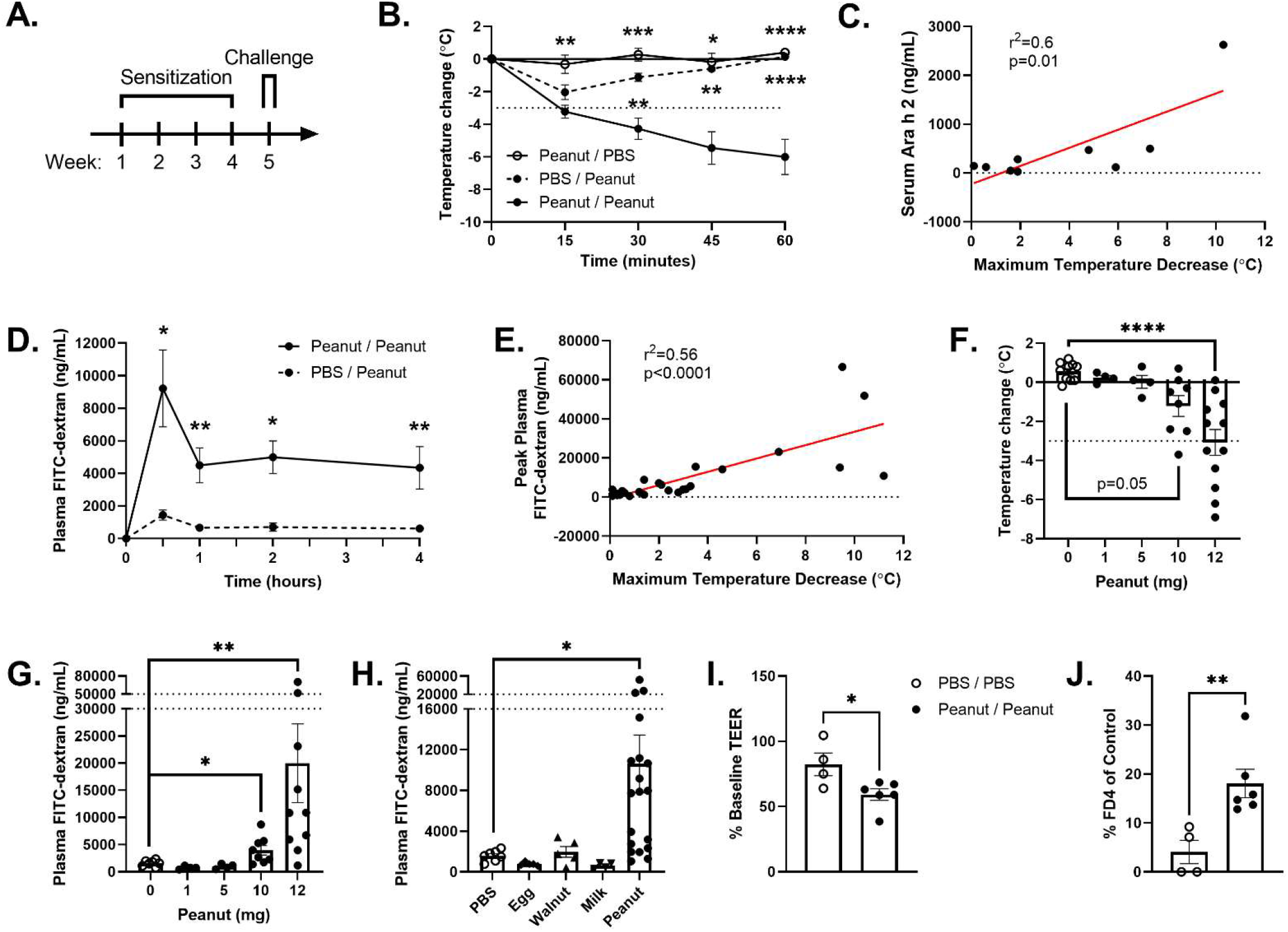
CC027 peanut allergy is associated with increased IEC permeability. **(A)** Mice received i.g. peanut once weekly for 4 weeks then were challenged Week 5. **(B)** Challenge rectal temperature change. CC027 PBS/Peanut: closed circle/dashed line (n=12); Peanut/PBS: open circle/solid line (n=8); Peanut/Peanut (closed circle/solid line, n=18). **(C)** Serum Ara h 2 thirty minutes after challenge versus maximum decrease in rectal temperature for each mouse (n=9), linear regression. **(D)** Plasma FD4 after challenge in PBS- (n=4) and peanut-sensitized (n=10) CC027 mice. **(E)** Peak plasma FD4 concentration versus maximum decrease in rectal temperature for each mouse (n=26), linear regression. **(F, G)** Peanut-sensitized CC027 mice were challenged with peanut (mg) i.g.: 0 (n=10), 1 (n=4), 5 (n=4), 10 (n=8), or 12 (n=12). Rectal temperature (F) and plasma FD4 (G) after 30 minutes. Kruskal-Wallis test. **(H)** Plasma FD4 thirty minutes after challenge with PBS (n=7), egg (n=5), walnut (n=5), milk (n=4), or peanut (n=19) in respectively orally-exposed CC027 mice. Kruskal-Wallis test, compared to PBS. **(I, J)** Jejunal EC isolated from PBS/PBS (n=4) and Peanut/Peanut (n=6) CC027/GeniUnc mice cultured on Transwell^®^ inserts. Transepithelial electrical resistance (TEER, I) and basolateral FD4 concentration (J) 15 minutes after addition of peanut to the apical compartment. Transit of FD4 across the IEC monolayer is expressed as the percentage of basolateral FD4 that moved across Transwells^®^ without cells present. Experimental duplicates were averaged. Multiple Mann-Whitney tests, compared to Peanut/Peanut.

Serum Ara h 2 thirty minutes post-challenge correlated with maximal decrease in rectal temperature in Peanut/Peanut CC027 (**Fig. 1C**) but not C3H (**Suppl. Fig. 1B**) mice. Proteins are primarily absorbed in the jejunum via transcellular transport (14), although alternative pathways may occur during pathology (7). Serum FITC-dextran 3-5 kDa (FD4) crosses the jejunal EC barrier paracellularly (15) and was administered i.g. alone or with peanut. Peanut/Peanut CC027 mice demonstrated elevated serum FD4 compared to that of PBS/Peanut CC027 mice at 30, 60, 120, and 240 minutes after challenge (**Fig. 1D**). This did not occur in Peanut/Peanut C3H mice (**Suppl. Fig. 1C**). Decreases in rectal temperature correlated with peak serum FD4 concentration in CC027 (**Fig. 1E**) but not C3H mice (**Suppl. Fig. 1D**). Rectal temperature and serum FD4 thirty minutes after challenge were significantly different between Peanut/PBS and Peanut/Peanut CC027 mice only with 10 mg and 12 mg (**Fig. 1F, G**). There was no difference in rectal temperatures (**Suppl. Fig. 1E**) or increase in plasma FD4 (**Suppl. Fig. 1F**) thirty minutes after challenge in C3H mice.

We similarly orally exposed mice to egg, walnut, or milk without adjuvant (13). CC027 mice demonstrated elevated serum egg- and walnut-, but not milk-, specific IgE, whereas C3H mice did not for any allergen (13). Thirty minutes post-challenge, only Walnut/Walnut CC027 mice demonstrated a significant but non-severe decrease in rectal temperature (13) but no increase in serum FD4 (**Fig. 1H**). C3H mice did not demonstrate rectal temperature change or increased plasma FD4 thirty minutes after challenge (**Suppl. Fig. 1G**). These data agree with detectable serum peanut allergen, but not milk or egg, after challenge (13). Eliciting doses of different allergens in an allergic individual are not known and are affected by both an individual’s intrinsic susceptibility and the food’s allergenicity (16). Non-protein components of foods may also play a role in allergenicity of a food (16). In this model, peanut was the most potent allergen, suggesting this model could be useful in understanding the differences between allergens and their effect on IEC barrier integrity.

A subtle defect in barrier integrity during IEC turnover (e.g., during IEC extrusion) could explain these results (17). A recent study using intravital confocal microscopy showed that patients with challenge-proven food allergy had increased ileal IEC barrier permeability due to defects during IEC extrusion at the villus tip (18); this can be used in the future to explore mechanisms of IEC turnover in the CC027 mouse.

A significant proportion of Peanut/Peanut CC027, but not C3H, mice developed watery diarrhea during challenge (**Suppl. Fig. 1H**) despite no differences in intestinal motility (**Suppl. Fig. 1I**). Increased IEC paracellular permeability was associated with secretory diarrhea in a food allergy model where i.p. OVA with alum-sensitized BALB/c mice received seven i.g. challenges with OVA 50 mg (19). Increased IEC paracellular permeability was also demonstrated in a peanut allergy model where C3H mice were sensitized by i.g. peanut with CT for six weeks prior to i.g. challenge (20). These studies found that intestinal mast cell (21) and goblet cell (20) numbers correlated with severity of reaction, but the effect of adjuvants on the IEC barrier in these models is not known. Secretory cell antigen passages (SAPs) provide rapid transit of antigen across the IEC barrier and delivery to subepithelial immune cells (22), which was necessary for an allergic reaction in one food allergy mouse model (23). Peanut/Peanut CC027 mice demonstrate more intestinal mast cells than similarly treated C3H and C57BL/6 mice (9), but SAPs have not been explored.

We used murine primary jejunal EC monolayer culture, in the absence of immune cells, to further explore IEC barrier integrity. Peanut/Peanut CC027-derived IECs had significant decreases in transepithelial electrical resistance (TEER) and increases in apical-to-basolateral FD4 flux when exposed to peanut (**Fig. 1I, J**). Although studies have associated increased IEC permeability with intestinal mast cell burden (21), this data suggest an immune cell-independent mechanism by which peanut affects IEC permeability. Notably, an initial immune cell-dependent process might be required to impart a durable change in IEC function and subsequent responses to allergens. Numerous immune cell products affect the IEC barrier (24), and mast cells from CC027 mice exhibit uniquely potent responses to intestinal parasite infection and passive cutaneous anaphylaxis (25). Additionally, IECs express the low-affinity Fcε receptor II (FcεRII, CD23), which shuttles IgE-antigen complexes transcellularly (26); this could explain the antigen-specific response of IECs when cultured in isolation from immune cells. Further studies will determine whether increased IEC barrier permeability in Peanut/Peanut CC027 mice is peanut-specific.

To investigate potential IEC-specific molecular contributors to allergen response, we isolated total RNA from jejunal IECs for RNA-seq at steady-state in peanut-sensitized mice and 30, 60, and 120 minutes after challenge (**Suppl. Fig. 2A**). Principal components analysis (PCA) revealed that sample transcriptomes had higher intra-experimental versus inter-experimental group similarity (**Fig. 2A**), suggesting distinct expression profiles at each timepoint. There were 13 differentially expressed genes (DEGs) in IEC after sensitization (FDR < 0.1), suggesting that sensitization itself does not significantly alter the jejunal EC expression profile under steady-state. We identified 183 (110 down- and 73 up-regulated), 266 (114 down- and 152 up-regulated), and 519 (217 down- and 303 up-regulated) DEGs 30-, 60-, and 120-minutes post-challenge, respectively (FDR < 0.1) (**Fig. 2B**). Gene enrichment analysis of the three time points showed late enrichment in cholesterol biosynthesis but few enriched pathways early in the time course (**Suppl. Fig. 2B**).

**Figure 2.**
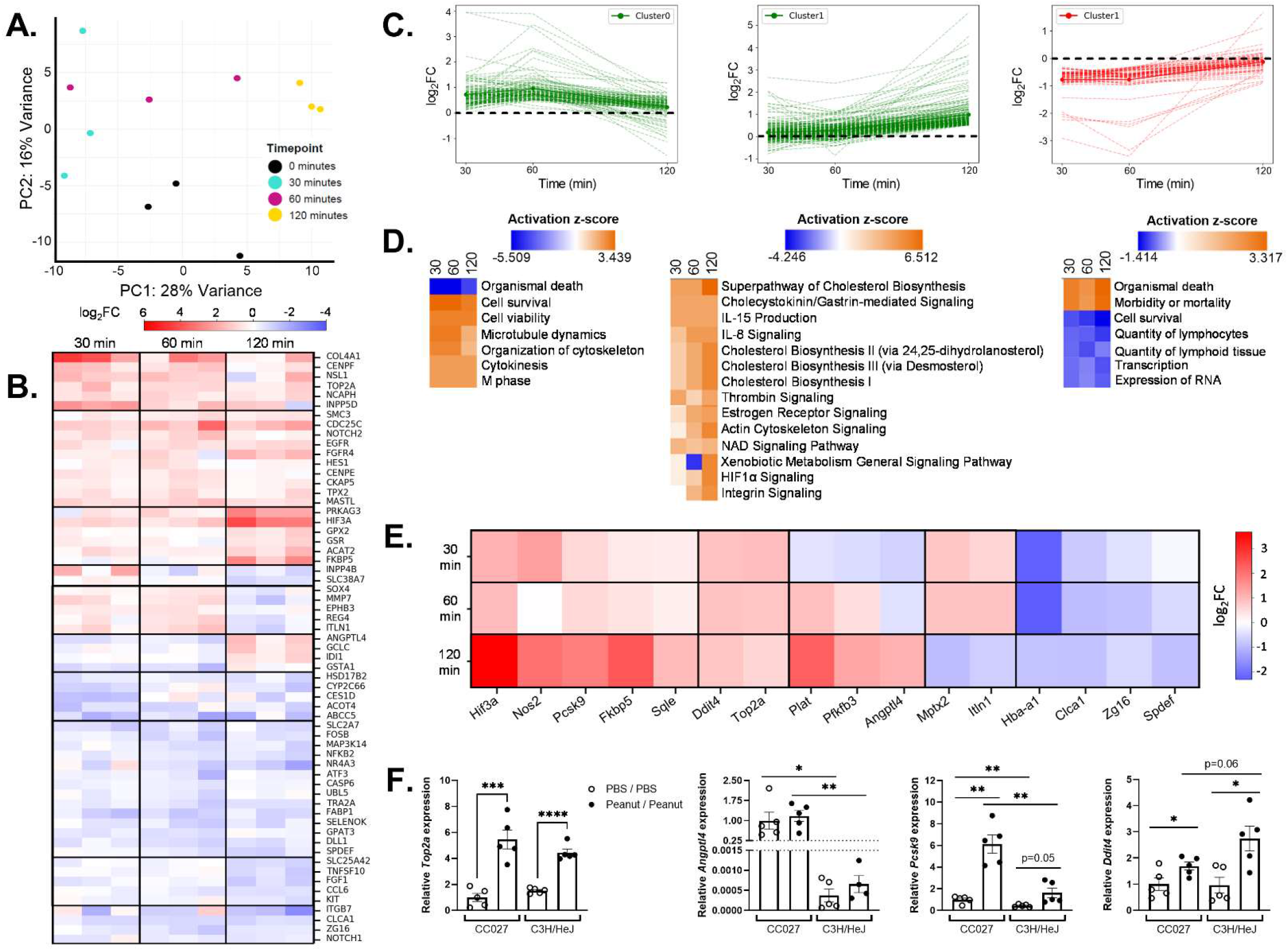
Jejunal EC transcriptomic response to peanut allergy. RNA-seq on jejunal EC isolated from PBS- or peanut-sensitized & unchallenged or challenged CC027/GeniUnc mice after 30, 60, or 120 minutes. n=3 per group. **(A)** PCA plot shows distinct expression profiles between jejunal EC from peanut-sensitized & unchallenged (black), or challenged mice after 30 (teal), 60 (magenta), or 120 (yellow) minutes. **(B)** Log_2_FC expression of genes from individual mice at each time point compared to peanut-sensitized & unchallenged CC027 mice. n=3 per group. **(C)** Expression profiles based on log_2_FC by time point of three clusters with >100 genes identified through time series analysis. **(D)** Enrichment analysis of genes in each cluster in (C). **(E)** Average differential expression (log_2_FC) at each timepoint of sixteen representative genes. n=3 per group. **(F)** Jejunal tissue expression by qRT-PCR by mouse strain and sensitization. n=5 per group. Welch’s t test.

To determine whether related genes showed similar variation in expression across timepoints, we generated a novel time series analysis by assigning a profile, based on the log_2_Fold-Change (log_2_FC) encompassing the three time points, to each of the 500 most DEGs and then performed cluster analysis (**Suppl. Fig. 2C**). We identified three clusters with >100 genes that showed similar changes in expression across timepoints (**Fig. 2C**) and performed gene enrichment analysis. Early and persistent programs of cell survival, proliferation and late induction of cholesterol biosynthesis pathways were enriched in our datasets (**Fig. 2D**). An early post-challenge increase in IEC survival and proliferation, perhaps due to dysregulated homeostasis, along with increased cholesterol biosynthesis to provide new cholesterol-rich cell membranes during proliferation (27), all support altered CC027 IEC homeostasis. Our RNA-seq analysis used only CC027 mice; future work will determine whether these findings are unique to this strain.

From the pathways enriched in our dataset, we identified 16 representative genes with the highest expression levels and log_2_FC with respect to the unchallenged mice (**Fig. 2E**). Quantitative RT-PCR from PBS/PBS and Peanut/Peanut CC027 and C3H mouse-derived jejunum 60 minutes after challenge showed different gene expression patterns by strain (**Fig. 2F, Suppl. Fig. 2D**), examples of which are shown in **Figure 2F**. For instance, DNA topoisomerase II alpha (*Top2a*) expression was increased in Peanut/Peanut CC027 and C3H mice. Angiopoietin-like 4 (*Angptl4*) showed significantly higher (~1000-fold) baseline and post-sensitization expression in CC027 compared to C3H. Proprotein convertase subtilisin/kexin type 9 (*Pcsk9*) baseline expression was higher in CC027 versus C3H jejuna, with greater increases after sensitization in CC027 mice. Conversely, DNA damage inducible transcript 4 (*Ddit4*) had higher baseline expression in CC027 jejuna with a greater increase after sensitization in C3H mice. The locations of four of the 16 genes (*Angptl4*, *Spdef*, *Fkbp5*, and *Plat*) lie within genomic regions where CC027/GeniUnc has highly divergent haplotypes compared to C3H. This and its 1000-fold higher expression in CC027 pointed to *Angptl4* as a putative relevant feature of peanut allergy in this model.

Angptl4 is a secreted glycoprotein regulated by microbiota, fasting, fatty acids, and hypoxia (12). It is proteolytically cleaved extracellularly into two products: the N-terminal coiled-coiled domain (nAngptl4) and C-terminal fibrinogen-like domain (cAngptl4). Full-length Angptl4 and nAngptl4 bind and inhibit lipoprotein lipase, altering lipid metabolism. *ANGPTL4* coding variants protect from cardiovascular disease (28). cAngptl4 may mediate vascular permeability, wound healing, and reactive oxygen species production (29). Angptl4 is primarily produced in the liver, adipose tissues, and to a lesser extent in the intestinal epithelium; its role in the intestines is poorly understood.

*Angptl4* expression by qRT-PCR was significantly higher in IECs versus lamina propria mononuclear cells in peanut-sensitized but unchallenged CC027 mice (**Fig. 3A**). Along the intestinal axis, the highest expression occurred in jejuna and lowest in colon from both PBS- and peanut-sensitized CC027 and C3H mice (**Fig. 3B, Suppl. Fig. 3A**). To determine whether altered ANGPTL4 translated to human peanut allergy patients, we explored intestinal *ANGPTL4* expression in healthy patients. RNA-seq data from healthy adults (30) showed that *ANGPTL4* expression is significantly higher in human ileal versus colonic tissue (**Fig. 3C**). Immunofluorescence in peanut-sensitized CC027 mice revealed Angptl4 protein was expressed in a subset of jejunal ECs (**Fig. 3D**), reflecting reported ANGPTL4 expression in human small IECs (https://www.proteinatlas.org/).

**Figure 3.**
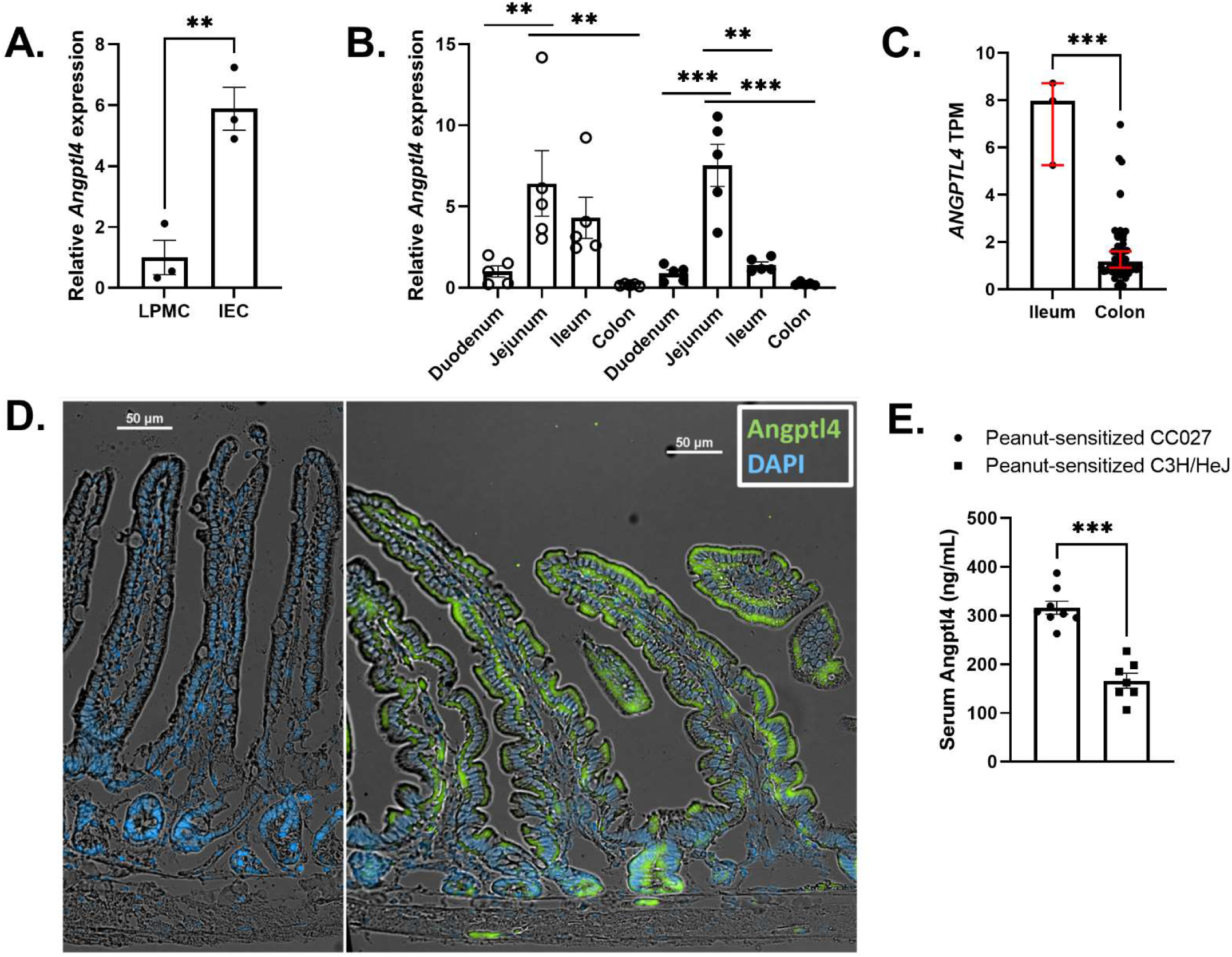
Murine and human Angptl4 expression is enriched in small intestinal epithelial cells. Angptl4 expression. **(A)** *Angptl4* expression in isolated IECs relative to lamina propria mononuclear cells (LPMC) from the same CC027 mouse by qRT-PCR. n=3 per group. Unpaired t test. **(B)** *Angptl4* expression in whole tissue from intestinal segments by qRT-PCR in PBS-sensitized (open circles) or peanut-sensitized (black circles) CC027 mice. n=5 per group. One-way ANOVA with multiple comparisons. **(C)** Normalized count by RNA-seq of *ANGPTL4* from ileal (n=3) or colonic (n=52) biopsies from healthy adult patients. Median with 95% CI. Mann-Whitney test. **(D)** IF of CC027 jejuna with anti-Angptl4 (green) and DAPI (blue) staining. Left panel is no 1° antibody. **(E)** Serum Angptl4 as measured by ELISA of peanut-sensitized CC027 (n=8) and C3H mice (n=7). Mann-Whitney test.

As ANGPTL4 is a secreted glycoprotein, we measured serum levels in peanut allergy. Serum Angptl4 was significantly higher in peanut-sensitized but unchallenged CC027 compared to C3H mice (**Fig. 3E**). Finally, serum ANGPTL4 was significantly higher in pediatric patients with challenge-confirmed peanut allergy compared to age-matched control patients without peanut allergy (**Table 1**). Control samples were from patients undergoing serum allergy testing and who had undetectable serum peanut-specific IgE. The control group included patients with allergy symptoms (though not peanut allergy), highlighting the potential use of ANGPTL4 as a specific biomarker of peanut allergy. Patient samples were collected at a separate date from the challenge visit and were therefore temporally unrelated to peanut exposure. We were still able to see a difference in serum ANGPTL4 between groups without controlling for metabolic state of the patient, which affects *ANGPTL4* expression (12). These data suggest that steady-state ANGPTL4 dysregulation accompanies peanut allergy. Future cohorts with comprehensive phenotyping of patients are needed to confirm these findings.

**Table 1.**
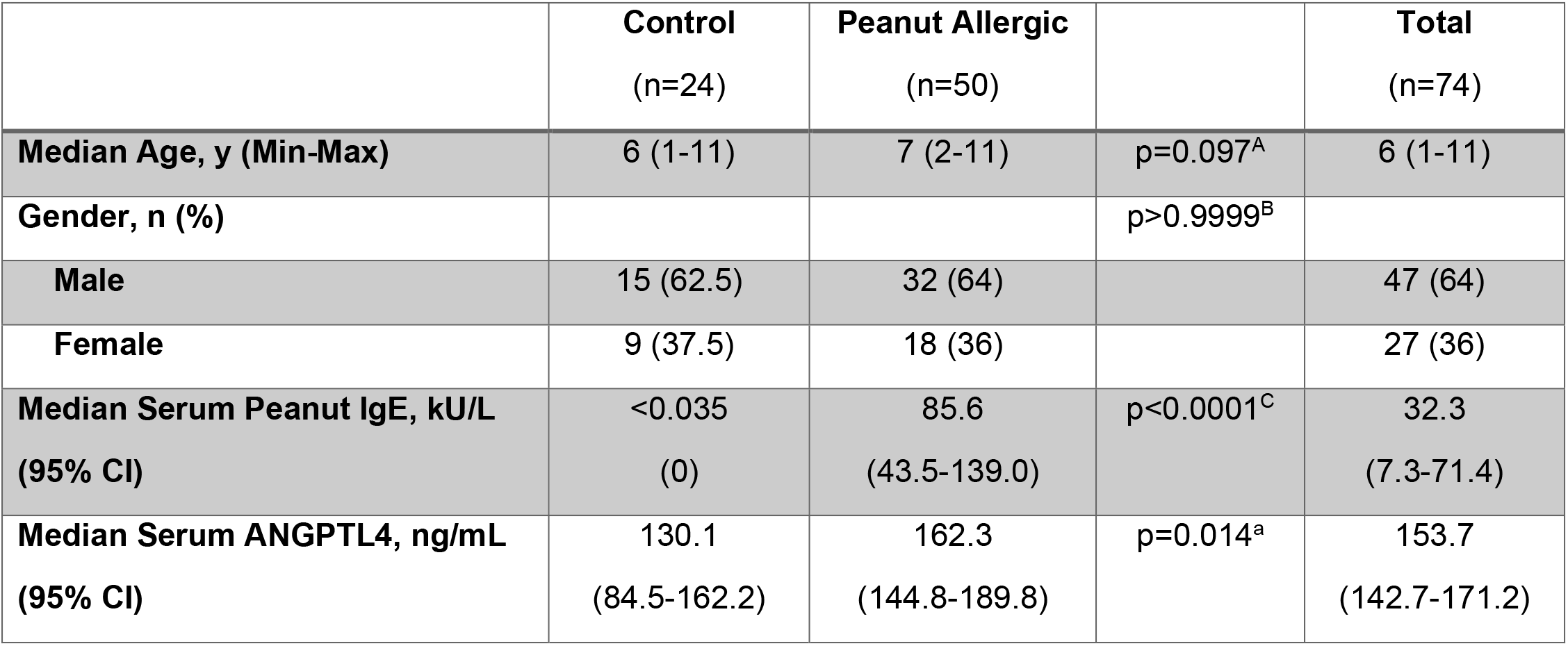
Peanut allergic pediatric patients have significantly elevated serum ANGPTL4 levels compared to non-peanut-allergic pediatric control patients. Food challenge-confirmed peanut allergic patients and age-matched non-peanut-allergic. Median age and gender proportion of the two groups were not significantly different. Median serum peanut-specific IgE was significantly higher in patients with peanut allergy. Median serum ANGPTL4 was significantly higher in patients with peanut allergy. ^A^Mann-Whitney test; ^B^Fisher’s exact test; ^C^One sample Wilcoxon test.

We report that IEC barrier integrity is disrupted in the adjuvant-independent CC027 model of peanut allergy, and IEC-predominant ANGPTL4 is revealed as a potential peanut allergy biomarker. These findings highlight the utility of the CC027 mouse model of peanut allergy to advance understanding of IEC dysfunction in human peanut allergy.

## Methods

Detailed methods are included in **Supplementary Materials**. All processed sequencing data are available in Gene Expression Omnibus under accession GSE179595.

### Study Approval

All procedures were approved under the UNC Institutional Review Board protocols 10-0355, 14-2445, 11-0359, and 11-2308. All animal experiments were approved by the UNC IACUC (#18-308).

## Author Contributions

E.C.S., S.Z.S. conceptualized the study; E.C.S, J.M.S., T.T., M.D.K., M.T.F., T.S.F., S.Z.S. designed the experiments; A.W.B., S.Z.S. supervised and funded the project; E.C.S., J.M.S., C.C., E.H.K. collected samples; E.C.S., J.M.S., A.B., L.P. performed experiments; E.C.S., J.M.S., S.R., T.T., L.P., M.D.K., T.S.F., S.Z.S. analyzed the data; E.C.S., S.Z.S. wrote the manuscript.

## Acknowledgments

This work was funded through Helmsley Trust, NIDDK P01DK094779, NIDDK 1R01DK104828-01A1, NIDDK P30-DK034987, NIAID U19AI100625, UNC CGIBD Training Grant (NIDDK, T32-DK007737), UNC Thurston Arthritis Research Center, UNC Physician-Scientist Training Program Fellowship, UNC/Duke University Allergy/Immunology Training Grant (NIAID, T32-AI007062), UNC SOM Office of Research, and NC TraCS Translational Team Science Award (TTSA017P1, TTSA017P2).

