## Supplementary Materials for "Increased intestinal permeability in an orally-reactive peanut allergy model identifies Angiopoietin like-4 as a biomarker"

**Supplementary Methods**

*Mice*

Female CC027/GeniUnc mice aged 4-6 weeks were obtained from the UNC Systems Genetics Core (1), C3H/HeJ mice were bred in house by MTF and provided in matched cohorts to the CC027 mice above. All mice upon receipt and rested for at least one week prior to starting experiments. CC027/GeniUnc mice were inbred for at least 20 generations to ensure 98% homozygosity. CC027/GeniUnc and C3H/HeJ strains were co-housed. Mice were kept on a 12:12 light:dark cycle and fed standard chow free of peanut, walnut, milk, and egg ingredients. All animal experiments were approved by the Institutional Animal Care and Use Committee at the University of North Carolina at Chapel Hill under protocol 18-308.

*Reagents*

| **Company** | **Reagent** | **Catalog Number** |
| --- | --- | --- |
| Golden Peanut (Alpharetta, GA) | Roasted and defatted peanut flour |  |
| Holmquist Hazelnut Orchards (Lynden, WA) | Roasted and defatted walnut flour |  |
| The Milky Whey (Missoula, MT) | Nonfat dry milk powder |  |
| Deb El Foods (Elizabethport, NJ) | Egg white powder |  |
| Sigma-Aldrich (St. Louis, MO) | Na_2_HPO_4_ | S7907 |
|  | KH_2_PO_4_ | P5655 |
|  | NaCl | S5886 |
|  | KCl | P5405 |
|  | FITC-conjugated dextran 3-5 kDa | FD4 |
|  | A83-01 | SML0788 |
|  | Accutase | A6964 |
| Thermo Fisher Scientific (Waltham, MA) | Sucrose | BP220-1 |
|  | D-sorbitol | BP439-500 |
|  | EDTA | 15575020 |
|  | Nicotinamide | AC128271000 |
|  | Gibco™ Recovery™ Cell Culture Freezing Medium | 12648010 |
|  | Serum-free B27™ 50X Supplement | 17504044 |
|  | Gibco™ HEPES buffer 1 M | 15630130 |
|  | Gibco™ GlutaMAX™ 100X Supplement | 35050061 |
|  | Applied Biosystems™ PowerUp™ SYBR™ Green Master Mix | 4367659 |
| Invitrogen™ (Thermo Fisher Scientific, Waltham, MA) | TRIzol™ Reagent | 15596026 |
|  | Rabbit ZO-1 Polyclonal Antibody | 61-7300 |
|  | Rabbit Occludin Polyclonal Antibody | 71-1500 |
|  | Goat anti-rabbit IgG Alexa Fluor 488 secondary antibody | A27034 |
|  | DAPI | D3571 |
|  | ProLong™ Gold Antifade Reagent mounting medium | P36934 |
| Worthington Biochemical (Lakewood, NJ) | Type IV Collagenase | LS004189 |
| PeproTech, Inc. (Rocky Hill, NJ) | Recombinant Murine EGF | 315-09 |
| APExBIO Technology, LLC (Houston, TX) | Y-27632 | A3008 |
| MP Biomedicals (Santa Ana, CA) | N-Acetyl-L-cysteine | 02194603-CF |
| InvivoGen (San Diego, CA) | Primocin™ | Ant-pm-05 |
| Anaspec (Fremont, CA) | Gastrin | AS-64149 |
| Selleck Chemical (Houston, TX) | SB202190 | S1077 |
| Cayman Chemicals (Ann Arbor, MI) | Prostaglandin E2 | 14010 |
| Proteintech® (Rosemont, IL) | Rabbit ANGPTL4 Polyclonal Antibody | 18374-1-AP |
| Norgen Biotek (Thorold, Ontario, Canada) | Total RNA Purification Plus Kit | 51800 |
| Sakura Finetec USA, Inc. (Torrance, CA) | O.C.T. (optimal cutting temperature) | 4583 |
| Biocare Medical (Concord, CA) | Reveal Decloaker solution | RV1000M |
| Dako (Glostrup, Denmark) | Protein Block Serum-Free | X0909 |
|  | Antibody Diluent | S0809 |
| LSBio, Inc. (Seattle, WA) | Human ANGPTL4 ELISA Kit | LS-F10823 |
|  | Mouse Angptl4 ELISA Kit | LS-F10824 |
| Indoor Biotechnologies (Charlottesville, VA) | Mouse Ara h 2 ELISA Kit | EPC-AH2-5 |

*Peanut, walnut, milk, and egg protein extractions*

Protein extractions were obtained as reported previously (2). Proteins were extracted from each of roasted and defatted peanut flour, roasted and defatted walnut flour, nonfat dry milk powder, or egg white powder in PBS (supplemented with 1 M NaCl for peanut and walnut extractions). Protein concentrations were measured by BCA (Pierce, Waltham, MA) and extracts were determined to contain all major allergens by SDS-PAGE gel.

*Sensitization and oral food challenges*

Mice were sensitized to food protein as previously described (3), with minor changes. Mice were fasted for at least 4 hours before and i.g. volume was 0.5 mL. Anaphylaxis was defined as >3°C decrease in rectal temperature.

*Measurement of IEC paracellular barrier permeability*

FD4 (600 mg/kg) was administered i.g. with peanut or PBS. Blood was collected by facial vein bleed into BD Microtainer® tubes containing lithium-heparin (BD Biosciences, Radnor PA). Samples were diluted 1:10-1:20 in PBS and fluorescence read on a ClarioStar plate reader. Values were normalized to that of mice given only PBS. Plasma FD4 concentration was determined using a FD4 standard curve (16 – 2000 ng/mL) diluted in normal mouse serum.

*Serum Ara h 2 and Angptl4 measurement*

Blood was collected by submandibular venipuncture at 30 or 60 minutes after challenge. Serum was diluted 1:10 and Ara h 2 was quantified via ELISA according to the manufacturer’s instructions (Indoor Biotechnologies, Charlottesville, VA). Serum was diluted 1:100 and Angptl4 was quantified via ELISA according to the manufacturer’s instructions (LS Bio, Inc., Seattle, WA). Sera from challenge-confirmed peanut allergic patients and non-peanut allergic patients were diluted 1:100 and ANGPTL4 was quantified via ELISA according to the manufacturer’s instructions (LS Bio, Inc., Seattle, WA).

*Tissue histology and immunofluorescence*

Freshly dissected intestines were sectioned into proximal (duodenum, from pyloric junction for 4 cm), middle (jejunum, starting 6 cm from pyloric junction for 4 cm), and distal (ileum, from cecal junction proximally for 4 cm) segments. One 2-cm piece each of duodenum, jejunum, ileum, and cecum, was cut open longitudinally on 4% paraformaldehyde (PFA)-soaked filter paper and submerged in 4% PFA at 4°C for at least 24 hours, washed in PBS, and cryoprotected first in 10% and then 30% sucrose, each overnight at 4°C. Frozen tissue embedded in O.C.T. was sectioned on a cryostat and slides stored at -80°C until use. Slides were thawed and washed in PBS. Antigen retrieval was performed in Reveal Decloaker solution in a Decloaking Chamber™ Pro (Biocare Medical, Concord, CA) with the following reaction protocol: 120°C for 30 seconds followed by 90°C for 10 seconds. Sections were permeabilized in 0.3% Triton-X in PBS, blocked with Blocking Solution, and stained with primary antibody in Antibody Diluent overnight at 4°C. After washing, tissue was incubated with secondary antibody in Antibody Diluent for 1 hour at room temperature. The sections were stained with DAPI, mounted with coverslips in ProLong™ Gold Antifade Reagent mounting medium, and visualized on a Nikon ECLIPSE Ti inverted microscope using NIS Elements v4.60.00 (Build 1171).

*IEC culture media*

| Medium | Reagent | Final Concentration |
| --- | --- | --- |
| Crypt Isolation Buffer (pH 7.4) | Na2HPO4 | 5.6 mM |
|  | KH2PO4 | 8.0 mM |
|  | NaCl | 96.2 mM |
|  | KCl | 1.6 mM |
|  | Sucrose | 43.4 mM |
|  | D-sorbitol | 54.9 mM |
| L-WRN Conditioned Medium | Generated as previously described (PMID: 28288348). |  |
| Complete Advanced DMEM/F12 Medium | Advanced DMEM/F12 Medium |  |
|  | Fetal Bovine Serum (FBS) | 20% v/v |
|  | Penicillin | 100 units/mL |
|  | Streptomycin | 100 μg/mL |
|  | GlutaMAX™ | 1X |
| Expansion Medium (EM) | L-WRN Conditioned Medium | 0.5X |
|  | Complete Advanced DMEM/F12 Medium | 0.5X |
|  | HEPES | 10 mM |
|  | GlutaMAX™ | 1X |
|  | Primocin™ | 100 μg/mL |
|  | N-Acetyl-L-cysteine | 1.25 mM |
|  | Nicotinamide | 10 mM |
|  | Gastrin | 10 nM |
|  | Recombinant murine EGF | 50 ng/mL |
|  | SB202190 | 3 μM |
|  | A83-01 | 500 nM |
|  | Prostaglandin E2 | 10 nM |
|  | B27 Supplement | 1X |
| Differentiation Medium (DM) | Complete Advanced DMEM/F12 Medium |  |
|  | HEPES | 10 mM |
|  | Primocin™ | 100 μg/mL |
|  | N-Acetyl-L-cysteine | 1.25 mM |
|  | Recombinant murine EGF | 50 ng/mL |
|  | A83-01 | 500 nM |

*Preparation of Gradient Collagen-Coated Transwell® Inserts*

Gradient collagen-coated Transwell® inserts for 12-well tissue culture plates were prepared as previously described (4).

*Isolation and culture of small intestinal epithelial cells*

Small intestinal crypts were isolated as previously described (5). Isolated crypts were either 1) frozen slowly overnight in Gibco™ Recovery™ Cell Culture Freezing Medium and stored in liquid nitrogen until future culture, 2) stored in TRIzol reagent at -80°C until RNA isolation, or 3) resuspended in Expansion Medium (EM) with 10 μM Y-27632 and grown on collagen-coated tissue culture plates. Freshly isolated IECs were resuspended in Expansion Medium (EM, 10^5^ cells/mL) and grown on gradient collagen-coated Transwell® inserts (37°C, 5% CO_2_). EM was replaced every 2 days, and for the first 2 days 10 μM Y-27632 was included. Monolayers were passaged on day 6-7. Monolayers and collagen were collected and collagen digested with 500 U/mL collagenase (Type IV, Worthington Biochemical LS004189, Lakewood, NJ) for 5 minutes at 37°C. After gentle pipetting to further dissociate the monolayers, centrifugation and removal of supernatant, cells were incubated with Accutase® in warmed PBS with 10 μM Y-27632 for 3 minutes at 37°C, washed, and resuspended in EM with 10 μM Y-27632 1:3 for continued culture.

For experiments, confluent monolayers were cultured in Differentiation Medium (DM) for at least 24 hours. Experiments were performed on days 4-6 when confluent monolayers were confirmed by transepithelial electrical resistance (TEER) >100 ohm × cm^2^, as measured by End-Ohm (World Precision Instruments, Inc.). Either 100 μg peanut extract or PBS and 10 mg FD4 were added to the apical compartment of the Transwell® insert and TEER measured every 5 minutes. Samples from the basolateral compartment were taken every 15 minutes for determination of FD4 concentration as described above.

*RNA isolation and RT-qPCR*

RNA isolation and quantitative RT-PCR was performed as previously described (5). RNA was extracted using TRIzol reagent and purified with the Total RNA Purification Plus Kit according to manufacturer’s instructions and concentration measured on NanoDrop™ spectrophotometer (Thermo Fisher Scientific, Waltham, MA). cDNA was made using 0.5 – 1.0 μg of RNA with the High-Capacity cDNA Reverse Transcription Kit (Applied Biosystems, Foster City, CA) on a thermocycler per manufacturer’s instructions. Samples were then diluted with nuclease-free water 1:5 and kept at -80°C until further use. One microliter of sample was used in one 20 μL PowerUp™ SYBR™ Green Master Mix reaction, and RT-PCR was run on QuantStudio 3 Real-Time PCR System (Applied Biosystems™, Waltham, MA). Samples were normalized to *Gapdh*.

*RT-qPCR Primers*

| **Gene ID (host)** | **Forward primer (5’🡪3’)** | **Reverse primer (5’🡪3’)** |
| --- | --- | --- |
| Spdef (mouse) | CACGTTGGATGAGCACTCGC | ACTTCTGCACGTTACCAGGGC |
| Angptl4 (mouse) | CAGAATCTTCAGAGCCAGATAGACC | GGAAAAGTCCACTGTGCCGC |
| Ddit4 (mouse) | TCCTCTTCGTCCTCGTCTCG | CCATCCAGGTATGAGGAGTCTTCC |
| Top2a (mouse) | TCGGGGCAAAAGAGTCATCCC | TAGTCTGCTTCTTTGCACCTGG |
| Mptx2 (mouse) | GTCCAAGAAGAGCACAATGGAG | AGGGCGAGTCAGGTCTGT |
| Hif3a (mouse) | AATGTCAGCAAGCACCTGGG | GCTTCTTCTTTGACAGGTTCGGC |
| Hba-a1 (mouse) | TGAAGCCCTGGAAAGGATGTTTG | AGCATCGGCGACCTTCTTGC |
| Zg16 (mouse) | TCGGCCTCTGCTAATTCCATTCAG | ACTGTGCCATAGCGCACCTG |
| Clca1 (mouse) | AGTATGGGCCACAAGACAGGAC | CCGGTAATGGCTGCTGAACAC |
| Itln1 (mouse) | ATAATGAGAGAGCGGCCAGTGC | CCATTGTGAGTTCCATATCCATCCC |
| Pcsk9 (mouse) | TATCCCAGCATGGCACCAGAC | GTCACACTTGCTCGCCTGTC |
| Fkbp5 (mouse) | TTTTGGAGAAGCCGGGAAGCC | GCGTGTACTTGCCTCCCTTG |
| Sqle (mouse) | CCCACAGTTACCTGAGCACCTG | TCCACCAGTAAGAGGGTGCC |
| Pfkfb3 (mouse) | GCTGACTCGCTACCTCAACTGG | AGCTAAGGCACACTGTTTTCGG |
| Nos2 (mouse) | TTGCCCCTGGAAGTTTCTCTTC | GGGATTCTGGAACATTCTGTGCTG |
| Plat (mouse) | AGCAAGCACTCTCGGGACAC | AGCCACGACTGGTGCTGTTG |
| Gapdh (mouse) | CGTCCCGTAGACAAAATGGT | TTGATGGCAACAATCTCCAC |

*RNA-seq processing and analysis*

Isolated jejunal ECs were stored in TRIzol at -80°C until RNA extraction. Library preparation and quality control testing were performed by Novogene (Sacramento, CA). Sequencing was performed on the Illumina NovaSeq platform using paired-end 150 bp reads. Data processing was performed as described previously (6). Transcript abundance was estimated by computing RPKM using RefSeq gene models aggregated by gene symbol. For differential expression analyses, raw counts over RefSeq exons were used, then compared across samples using DESeq2 (7) at an FDR of 10%.

*Time Series Analysis*

We arranged differentially expressed genes in increasing order of adjusted p-value (*padj*) from three time-points: 30; 60; 120; and created separate charts for up- and down-regulated genes. After selecting the top *k* = 500 genes from each list, each gene was represented as a profile of its *log2FC* in the three timepoints, i.e., $v\left( g \right)=\left[ l_{30}^{g} , l_{60}^{g} , l_{120}^{g} \right]$*,* where $l_{t}^{g}$is the *log2FC* of gene $g$ at time-point $t$. We calculated the similarity in differential expression profiles between two genes $g_{i}$ and $g_{j}$ in terms of correlation between their gene profiles $v\left( g_{i} \right)$ and $v\left( g_{j} \right)$, i.e., $\left| PCC \left( v\left( g_{i} \right), v\left( g_{j} \right) \right) \right|$. We created a network *G*, where the genes are represented as nodes ($g\in V$) and an undirected link $\left( g_{i}, g_{j} \right)$exists between each pair of genes $g_{i}$ and $g_{j}$. Each undirected link $\left( g_{i}, g_{j} \right)$ may exist (and have a weight $w_{i,j}$) depending on the network variant (discussed hereafter). Depending on the mode of the *log2FC* and Pearson correlation coefficient, we can create variants of network *G*. We specifically considered the following network variants:

- Fold change. We constructed network *G* comprising genes that are either (1) only positively expressed (abbreviated **p**) or (2) only negatively expressed (abbreviated **n**) across all three time points 30, 60, 120.
- Nature of correlation. Pearson correlation coefficient can be positive or negative. We consider three types here:

1. Positive correlation only (abbreviated +1). Undirected link $\left( g_{i}, g_{j} \right)$ exists in *G* with weight $w_{i,j}$ = $\left| PCC \left( v\left( g_{i} \right), v\left( g_{j} \right) \right) \right|$ if and only if $PCC \left( v\left( g_{i} \right), v\left( g_{j} \right) \right)>0$.
2. Negative correlation only (abbreviated -1). Undirected link $\left( g_{i}, g_{j} \right)$ exists in *G* with weight $w_{i,j}$ = $\left| PCC \left( v\left( g_{i} \right), v\left( g_{j} \right) \right) \right|$ if and only if $PCC \left( v\left( g_{i} \right), v\left( g_{j} \right) \right)<0$.
3. Sign agnostic or unsigned correlation only (abbreviated 0). Undirected link $\left( g_{i}, g_{j} \right)$ exists in *G* with weight $w_{i,j}$ = $\left| PCC \left( v\left( g_{i} \right), v\left( g_{j} \right) \right) \right|$, irrespective of sign of the correlation.

We applied agglomerative hierarchical clustering on similarity network *G* to iteratively group gene vectors with highly (positive or negative) correlated gene vectors, resulting in the formation of clustered network structures. We measured cluster quality using the *Calinski and Harabasz score*, which is the ratio between inter- and intra-cluster dispersion. The score is higher when clusters are dense and well separated, which relates to a standard concept of a cluster (8,9). Finally, the important genes in each cluster were identified using network centrality measures. In each cluster *c* with gene set $g_{j}\in Z_{c}$, we measured degree centrality of gene $g_{j}$as the sum of weights of links connected to $g_{j}$ in cluster *c*, i.e., $\sum_{g_{j}\in Z_{c}} w_{i,j}.$

*Human RNA-seq Data*

Processed RNA-seq data from ileal and colonic biopsy tissue from healthy non-IBD patients were previously published (6). All processed sequencing data are available in Gene Expression Omnibus (GEO) under accession GSE85499 with raw sequence data available through dbGaP.

*Statistical methods*

Data is shown as mean ± SEM unless otherwise noted. Except for RNA-seq data, all statistical analyses were performed using Prism (GraphPad Software). Exact tests used are specified in the figure legends. Statistical significance is defined as *p< 0.05, **p< 0.01, ***p< 0.001, ****p<0.0001.

**Supplementary Figures**

**Supplementary Figure 1. Peanut exposure in C3H/HeJ mice is not associated with increased IEC permeability.** **(A)** Rectal temperature in peanut-exposed & PBS-challenged (Peanut/PBS; open circle/solid line, n=5), and peanut-exposed & -challenged (Peanut/Peanut; closed circle/solid line, n=10) C3H/HeJ mice after challenge with i.g. 12 mg peanut. Multiple Mann-Whitney tests, compared to Peanut/Peanut. **(B)** Serum Ara h 2 thirty minutes after challenge measured by ELISA were plotted against the maximum decrease in rectal temperature for each mouse (n=9), and linear regression performed. **(C)** PBS- (n=5) and peanut-exposed (n=10) C3H/HeJ mice were given i.g. 12 mg peanut with 600 mg/kg FD4, and plasma was collected. Plasma FD4 concentration was determined with a standard curve. Multiple Mann-Whitney tests, compared to Peanut/Peanut. **(D)** Peak plasma FD4 concentration versus the maximum decrease in rectal temperature for each mouse (n=7); linear regression. **(E, F)** Peanut-exposed C3H/HeJ mice were challenged with i.g 0 (n=3), 1 (n=4), 5 (n=4), 10 (n=4), or 12 mg (n=4) peanut, and rectal temperature (E) and plasma FD4 concentrations (F) were measured after 30 minutes. Kruskal-Wallis test. **(G)** PBS- (n=4), egg- (n=4), walnut- (n=4), milk- (n=3), and peanut-exposed (n=4) C3H/HeJ mice were challenged i.g. with respective foods, and plasma FD4 concentration determined after 30 minutes. Kruskal-Wallis test, compared to PBS. **(H)** Proportion of Peanut/Peanut mice by strain with watery diarrhea during challenge. Fisher’s exact t test. **(I)** Small intestinal motility as measured by percent of small intestine length that carmine dye/cellulose solution traversed 20 minutes after i.g. administration by sensitization in CC027/GeniUnc (circles) and C3H/HeJ (squares) mice. n=8 per group. Multiple Mann-Whitney tests.

**Supplementary Figure 2. CC027/GeniUnc jejunal EC expression profiles highlight response to peanut allergy. (A)** RNA-seq was performed on jejunal EC isolated from PBS- or peanut-sensitized & unchallenged or challenged CC027/GeniUnc mice after 30, 60, or 120 minutes. n=3 for each group. **(B)** Gene enrichment analysis (Ingenuity Pathway Analysis, Qiagen) showed strong upregulation of cholesterol biosynthesis pathways late after challenge. **(C)** Time series analysis was performed by assigning expression profiles to each differentially expressed gene over the three time points then clustering based on similarity. **(D)** Whole jejunal tissue expression by qRT-PCR by mouse strain and sensitization. n=5 for each group. Welch’s t test.

**Supplementary Figure 3. Murine Angptl4 expression is enriched in small intestine of C3H/HeJ mice.** Angptl4 expression was measured by qRT-PCR. **(B)** *Angptl4* expression in whole tissue from intestinal segments in PBS- (open squares) or peanut-exposed (black squares) C3H/HeJ mice. n=5 per group. One-way ANOVA with multiple comparisons.

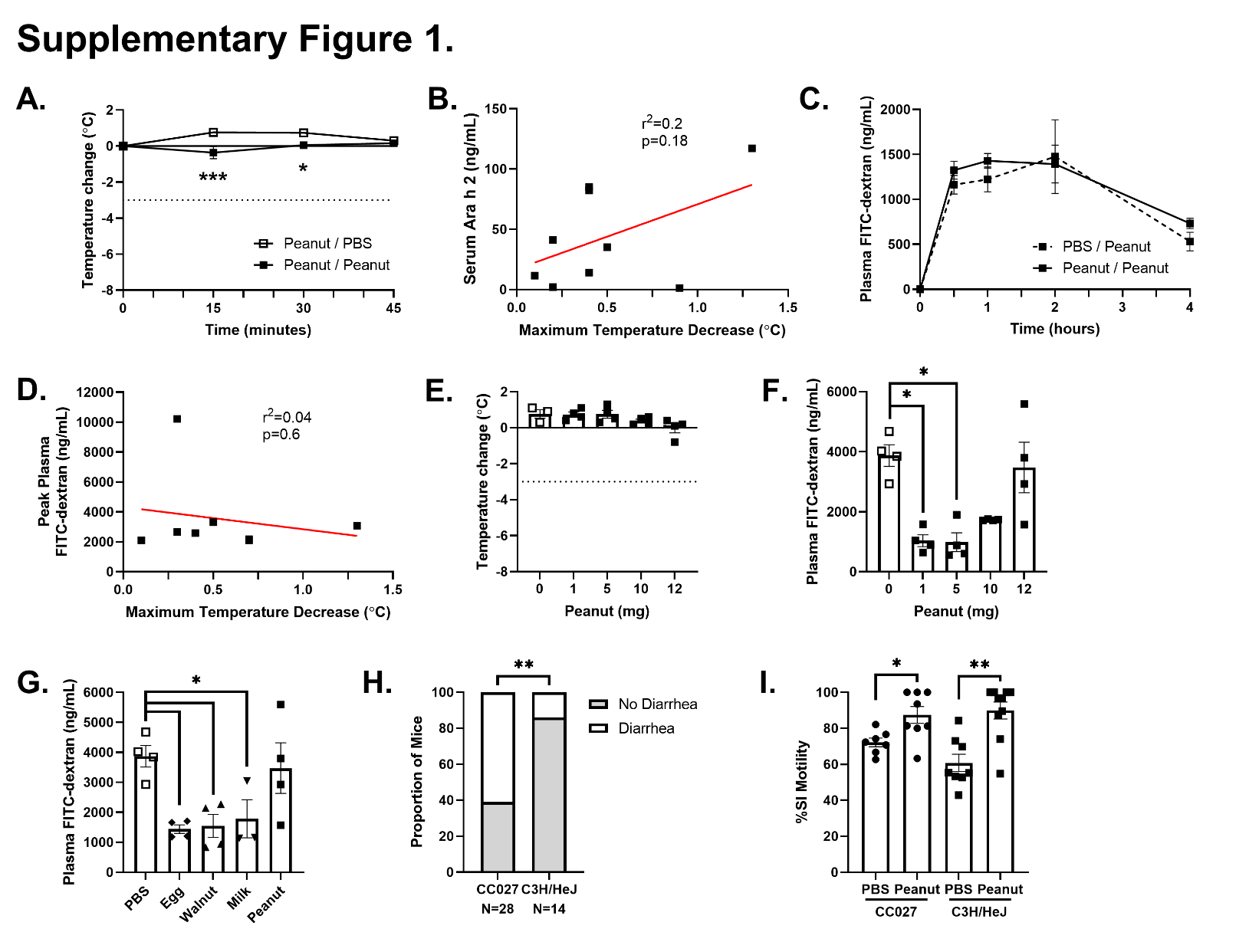

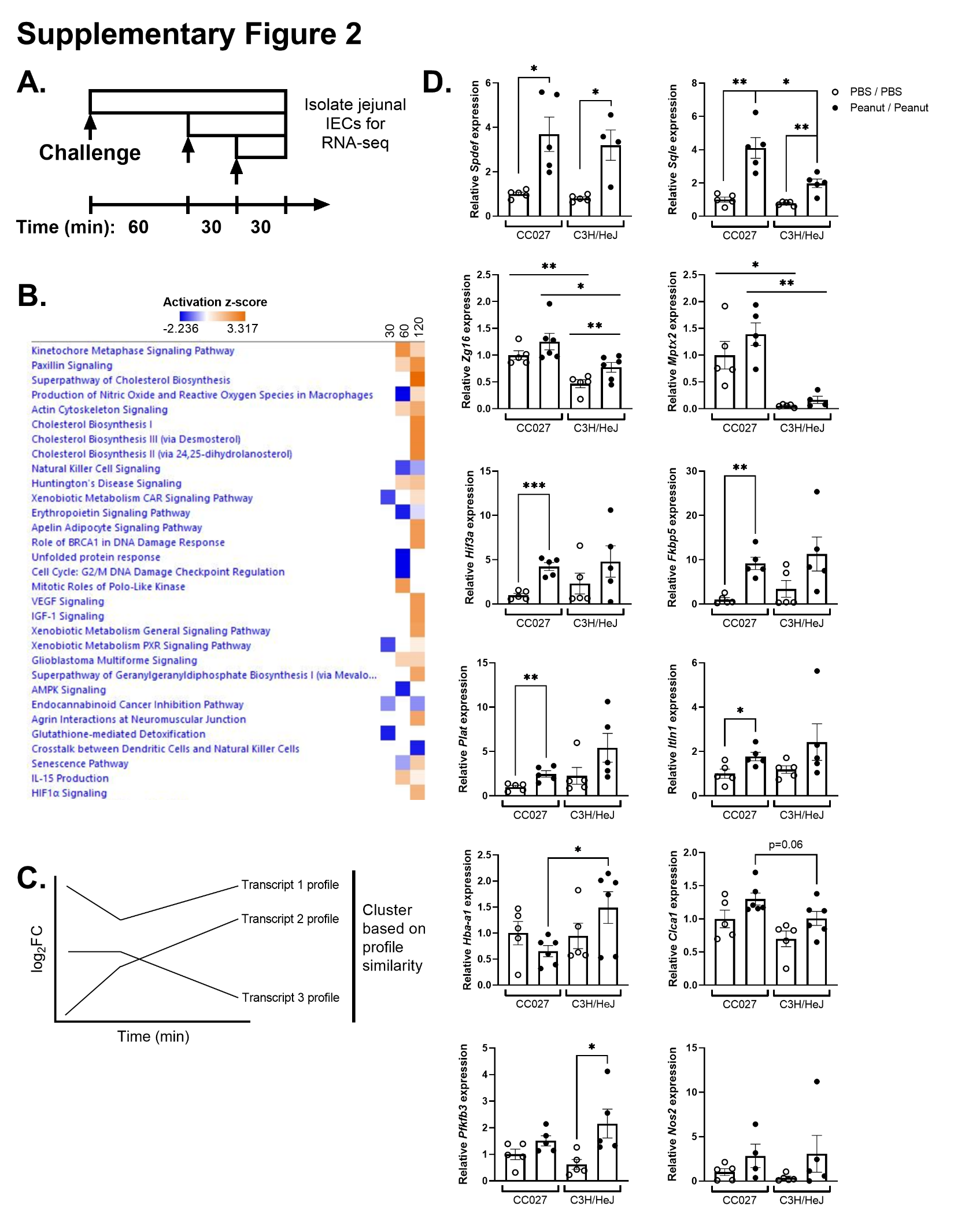

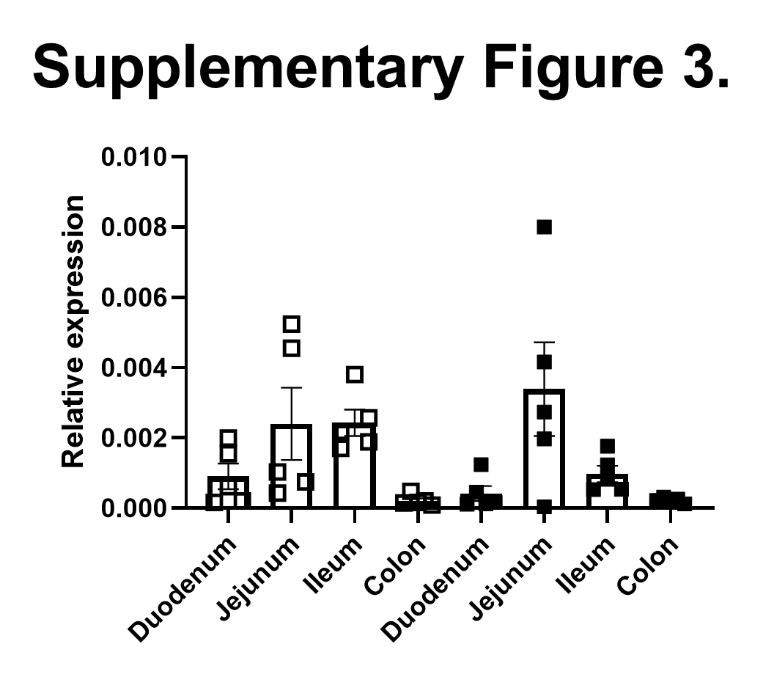
